# The evolution of thermal performance in native and invasive populations of *Mimulus guttatus*

**DOI:** 10.1101/2020.09.10.291252

**Authors:** Aleah Querns, Rachel Wooliver, Mario Vallejo-Marín, Seema Nayan Sheth

## Abstract

**1.** The rise of globalization has spread organisms beyond their natural range, allowing further opportunity for species to adapt to novel environments and potentially become invaders. Yet, the role of climatic niche evolution in promoting the success of invasive species remains poorly understood. Here, we use thermal performance curves (TPCs) to test hypotheses about thermal adaptation during the invasion process. First, if novel temperature regimes impose strong selection in the introduced range, invasive populations may evolve distinct TPCs relative to native populations. Second, invasive species may not exhibit specialist-generalist tradeoffs and instead may be “masters-of-all” with high maximum performance and broad TPCs. Third, with sufficient time, standing genetic variation, and temperature-mediated selection, TPCs of native and invasive populations may show parallel evolution in response to thermal gradients.
**2.** To test these hypotheses, we built TPCs for 18 native (United States) and 13 invasive (United Kingdom) populations of the yellow monkeyflower, *Mimulus guttatus*. We grew clones of multiple genotypes per population across six temperature regimes in growth chambers.
**3.** Invasive populations have not evolved different thermal optima or performance breadths, providing evidence for evolutionary stasis of thermal performance between the native and invasive ranges after over 200 years post-introduction. Further, both native and invasive populations experienced similar specialist-generalist tradeoffs whereby broad TPCS come at the cost of lower peak performance. Inconsistent with the idea that the degree of thermal specialization varies across spatial or temperature gradients, native and invasive populations did not exhibit adaptive clines in thermal performance breadth with latitude or temperature seasonality. However, thermal optimum increased with mean annual temperature in the native range, indicating some adaptive differentiation among native populations
**4.** *Synthesis:* These findings suggest that thermal niches were static during the invasion process, and that general-purpose genotypes, rather than rapid evolution in the introduced range, may promote invasion.

## Introduction

Invasive species are one of the greatest threats to biodiversity in the 21st century, and while the importance of managing their spread is widely recognized, the factors that contribute to a species becoming invasive are still debated (Sala et al., 2000; Lee, 2002; van Kleunen, Dawson, et al., 2015). Baker’s classic description of “ideal weeds” suggested that invasive species originate from “general-purpose” genotypes with broad climatic tolerance (Baker, 1965). However, it is unknown whether climatic tolerance in a species invasive range is pre-existing, which predicts niche conservatism, or evolved, which predicts niche lability between the native and invasive range (Petitpierre et al., 2012; Atwater et al., 2018; Liu et al., 2020). Upon introduction to a new geographic region, a species may expand, contract, or maintain its climatic niche space depending on how the set of environmental conditions required to support non-negative population growth changes from the native to the invasive range (Broennimann et al., 2012). Similarly, a species average niche conditions (often referred to as niche “centroid” or “position”) may or may not shift in the invasive range (Guisan et al., 2014). If climatic niches are often conserved between species native and invasive ranges, then building climatic niche models based on native occurrences to identify areas with high invasion risk holds great promise for predicting and managing biological invasions (Thuiller et al., 2005; Chapman et al., 2017; Da Re et al., 2020). Numerous studies have used correlative niche models to assess climatic niche conservatism across species native and invasive ranges (Peterson et al., 2003; Kriticos et al., 2013; Da Re et al., 2020), but most have neglected among-population variation in niche properties and variation in physiological tolerances across climatic gradients resulting from evolutionary change (some exceptions include Ebeling et al., 2008; Hill et al., 2013). We use an experimental approach to evaluate the evolution of physiological tolerances within and between a species native and invasive ranges.

One dimension of a species climatic niche that influences its ability to invade new areas is temperature. Thermal performance curves (TPCs, Fig. 1A) describe the performance of a genotype, individual, population, or species across a temperature gradient (Huey & Stevenson, 1979). Though TPCs are not strictly equivalent to thermal niches unless the performance metric is total fitness, they provide a powerful means of experimentally approximating a species fundamental thermal niche. Comparisons of TPCs may not explicitly test thermal niche conservatism between a species native and introduced ranges, but they shed light on whether the evolution of physiological tolerance plays a role in species abilities to invade new areas. Some TPC parameters that have been hypothesized to influence adaptation to temperature gradients include maximum performance, thermal optimum, thermal performance breadth, and area under the curve (Huey & Stevenson, 1979; Angert et al., 2011; Fig. 1A). Maximum performance is the highest level of performance achievable, corresponding to the peak of the TPC. Thermal optimum details the temperature at which maximum performance is achieved (Huey & Stevenson, 1979), analogous to thermal niche position (or centroid). Comparable to thermal niche breadth, thermal performance breadth describes the span of temperatures across which a specified percentage of the maximum performance is achieved (Huey & Stevenson, 1979), with specialists having narrow breadth, and generalists having wide breadth. Finally, the area under the curve refers to the total amount of area between a TPC and the temperature axis, analogous to total thermal niche space.

**Figure 1.**
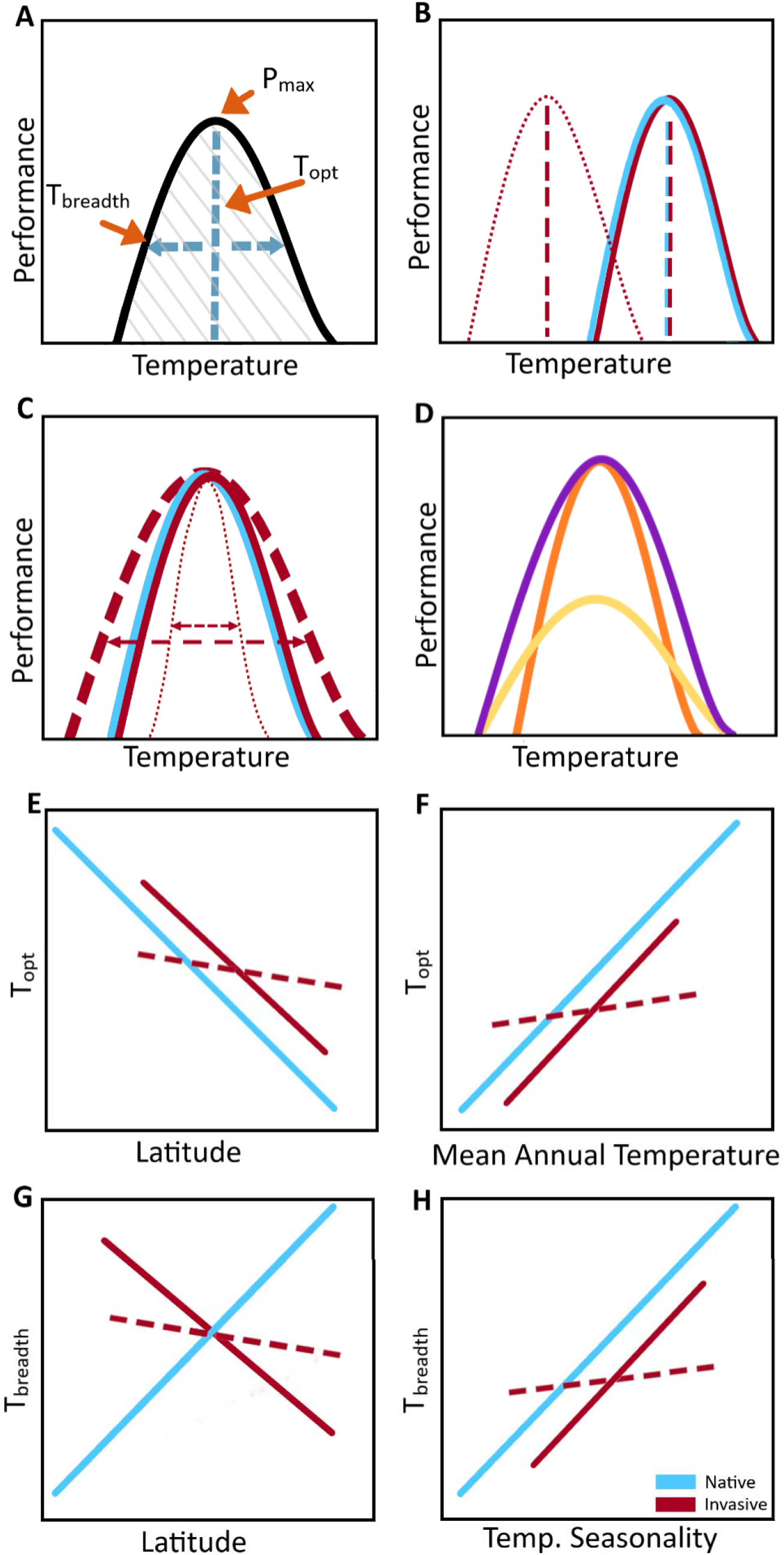
Hypotheses describing the evolution of thermal performance curves (TPCs) and clines in native and invasive populations of *M. guttatus*. (A) TPC parameters of interest include maximum performance (*P_max_*), thermal optimum (*T_opt_*), thermal performance breadth (*T_breadth_*), and area under the curve (shaded region) (B) Invasive populations may exhibit a lower *T_opt_* relative to native populations (dotted line) due to lower average mean annual temperatures (MAT; Fig. 2C; native average MAT= 9.2°C, invasive average MAT=8.6°C). Alternatively, native and invasive populations may exhibit similar *T_opt_* if populations occupy similar thermal regimes or if *T_breadth_* is maintained (solid lines) in the invasive range. (C) Invasive populations may evolve wider (dashed line) or narrower (dotted line) *T_breadth_* relative to native populations due to admixture or bottlenecks/reduced temperature variation, respectively. Alternatively, if general-purpose genotypes from native populations facilitate invasion, *T_breadth_* may be maintained (solid line). (D) Within native and invasive populations, there may be specialist-generalist tradeoffs such that populations with greater *T_breadth_* have a lower *P_max_* (yellow) and populations with narrower *T_breadth_* have a higher *P_max_* (orange). However, some invasive species may not exhibit these tradeoffs and instead achieve a wide *T_breadth_* and a high *P_max_* (purple). (E-F) The native range (solid blue line) may exhibit clines such that *T_opt_* decreases with latitude (E) and increases with MAT (F), and these clines may be parallel (solid red) or weaker/nonexistent (dashed red) in the invasive range. (G-H) The native range may exhibit clines such that *T_breadth_* increases with latitude (G) and temperature seasonality (H). The invasive range, where temperature seasonality increases with latitude (Fig. 2D), may evolve parallel clines (solid line) such that *T_breadth_* decreases with latitude (G) and increases with temperature seasonality (H), or may exhibit weaker/no clines (dashed line).

Evolutionary divergence of TPC parameters between invasive and native populations of the same species could arise through thermal niche contraction, expansion, and/or shifts in the introduced range. Consistent with thermal niche shifts, changes in thermal optima may accompany divergence in local thermal regimes between the native and invasive range (Angiletta, 2009). Nonetheless, if populations possess broad thermal performance breadths, and/or if temperature regimes are similar in both ranges, then native and invasive populations may exhibit similar thermal optima (Guisan et al., 2014). Consistent with niche expansion, invasive populations could evolve a broader TPC upon introduction to novel thermal regimes (Zerebecki & Sorte, 2011; Bates et al., 2013). Populations can achieve broad TPCs through a combination of phenotypic plasticity and standing genetic variation (Sheth & Angert, 2014), particularly if there is admixture of multiple source populations introduced to the novel range (Kolbe et al., 2004; Lavergne & Molofsky, 2007). Though genetic bottlenecks during founding events may constrain the ability of invasive populations to adapt to novel conditions, or cause reductions in breadth (consistent with thermal niche contraction) due to genetic drift (Kitayama & Mueller◻Dombois, 1995; Daehler, 2003), invasive populations rarely undergo significant reductions of genetic diversity relative to native populations, making this scenario unlikely (Wares et al., 2005; Dlugosch & Parker, 2008). Alternatively, if source populations represent general-purpose genotypes, they may be predisposed toward having a broad thermal niche which may be conserved in the invasive range (Ainsworth & Drake, 2020), leading to similar thermal performance breadths in native and invasive populations. To date, there are few empirical studies which compare TPCs of native and invasive populations of a single species (but see Comeault et al., 2020), particularly in plants.

Evolutionary trajectories in the introduced range may be influenced by energetic tradeoffs in thermal reaction norms (Angilletta et al., 2003). Commonly referred to as “jack-of-all-trades is a master of none” and specialist-generalist tradeoffs, the ability to persist over a wider range of temperatures can come at the cost of lower maximum performance (Huey & Hertz, 1984; Richards et al., 2006; Le Vinh Thuy et al., 2016). Such tradeoffs should result in a negative relationship between maximum performance and thermal performance breadth across populations, whereby populations with broader TPCs have lower peak performance due to additional energetic costs (Dewitt et al., 1998; Angilletta et al., 2003; Richards et al., 2006). This tradeoff operates under the assumption that the area under the TPC remains constant (Palaima & Spitze, 2004), but physiological or biochemical alterations could allow higher performance across a broad range of temperatures (Angilletta et al., 2003; Richards et al., 2006) and cause a positive relationship between thermal performance breadth and maximum performance. This phenomenon, which has been referred to as “jack-and-master”, or “master of all” (Richards et al., 2006; Matesanz & Sultan, 2013), may be one mechanism that advances invasion.

Given a temperature or latitudinal gradient, populations with sufficient time and genetic variation may evolve clines in TPC parameters in response to temperature-mediated selection (Endler, 1977; Diamond et al., 2017; Campbell◻Staton et al., 2018). Failure to account for the presence of latitudinal or environmental clines can mask inferences of trait evolution between native and invasive ranges, as divergent selection may occur among populations (Colautti et al., 2009). Adaptation to local thermal regimes should result in thermal optimum increasing with mean annual temperature (Angert et al., 2011), and thermal performance breadth increasing with temperature seasonality (often referred to as the climate variability hypothesis, CVH; Dobzhansky, 1950; Janzen, 1967; Stevens, 1989; Gutiérrez-Pesquera et al., 2016). Thus, parallel phenotypic clines across latitudinal or climatic gradients in native and invasive ranges would indicate that rapid evolution plays a role in the invasion process (Huey et al., 2000; Hernández et al., 2019; van Boheemen et al., 2019). However, if invasive populations have not had sufficient time or standing genetic variation for adaptation to novel temperature gradients, or if temperature-mediated selection in the introduced range is weak, then phenotypic clines may be shallower in the invasive range than in the native range, or absent altogether (Bhattarai et al., 2017).

In this study, we compare TPCs of invasive and native perennial populations of the yellow monkeyflower, *Mimulus guttatus* (Phrymaceae; Fig. 2A,B), also known as *Erythranthe guttata* (Fraga, 2018; Lowry et al., 2019), to improve our understanding of species invasive abilities. *Mimulus guttatus* is an herbaceous plant native to western North America, occupying wet habitats across a broad latitudinal and climatic gradient from Alaska to Northern Mexico (Fraga, 2018). In 1812, *M. guttatus* was brought to the United Kingdom (UK) for horticultural use and has subsequently become widespread across the British Isles and to a lesser extent parts of continental Europe (Preston et al., 2002; Newman, 2015). *Mimulus guttatus* is now considered invasive in the UK and New Zealand (Vallejo-Marín & Lye, 2013; van Kleunen, Röckle, et al., 2015; Fraga, 2018). Genomic data suggest that genotypes of *M. guttatus* in the UK originated from perennial populations in Alaska (Puzey & Vallejo-Marín, 2014; Pantoja et al., 2017), followed by multiple introductions of perennial populations from across the native range (Vallejo-Marín et al., 2020). The invasive UK range thus consists of a highly admixed melting pot of native populations and is not likely constrained by low genetic diversity. Climatic niche models indicate niche conservatism in the native and invasive European ranges of *M. guttatus*, though invasive populations occur in a subset of conditions occupied in the native range (Da Re et al., 2020). Because the non-native populations are locally dominant invaders in the UK compared to continental Europe, and served as a bridgehead for invasions worldwide (Vallejo-Marín et al., 2020), we focused exclusively on the UK portion of the invasive range.

**Figure 2.**
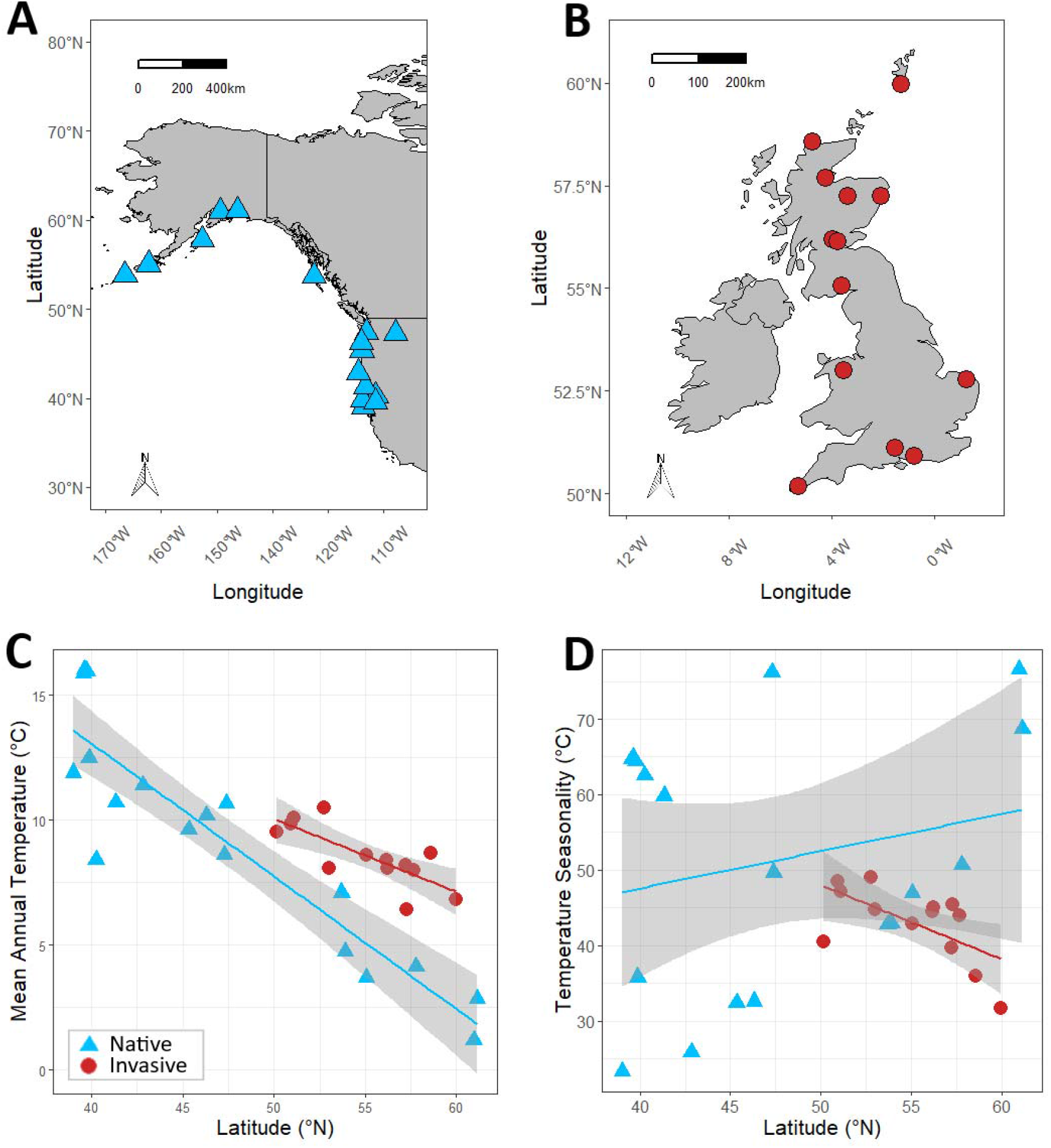
Map of focal populations of *M. guttatus* in (A) native range in North America and (B) invasive range in the United Kingdom. Relationships between (C) mean annual temperature and (D) temperature seasonality with latitude for native and invasive populations. Fitted lines indicate predicted mean annual temperature or seasonality as a function of latitude based on linear models. The gray shaded area represents a 95% confidence interval for these predictions. Climate data were obtained from WorldClim v.2 (~1-km resolution, 1970-2000, Fick & Hijmans, 2017).

Here, we assess hypotheses about whether thermal performance evolution and the ability to overcome specialist-generalist tradeoffs contribute to the invasion success of *M. guttatus*. First, we hypothesized that if the species has undergone changes in thermal regimes and/or genetic variation in its invasive range, there would be changes in thermal optimum and performance breadth between the native and invasive ranges, against the alternative hypothesis that these parameters do not differ between ranges (Fig. 1B,C). Since average temperatures are slightly higher in the native range (Fig. 2C), invasive populations may have evolved lower thermal optima relative to native populations (Fig. 1B). Further, while admixture in the invasive range could lead to broader TPCs relative to the native range, reduced temperature variation (Fig. 2C,D) and/or genetic bottlenecks could result in narrower TPCs in the invasive range (Fig. 1C). Second, we evaluated the hypothesis that the ability to overcome specialist-generalist tradeoffs has fostered the invasion of *M. guttatus* (Fig. 1D) such that native and invasive populations are “masters-of-all”. Alternatively, populations may exhibit specialist-generalist tradeoffs, predicting that “jack-of-all-trades is a master of none”. Third, given sufficient time, standing genetic variation, and temperature-mediated selection, we hypothesized parallel evolution in response to thermal gradients in the invasive range (Fig. 1E-H). We predicted that thermal optimum would increase with mean annual temperature (Fig. 1F), which decreases with latitude in both ranges (Fig. 1E; Fig. 2C), and thermal performance breadth would increase with temperature seasonality (Fig. 1H), which increases with latitude in the native range but decreases with latitude in the invasive range (Fig. 1G; Fig. 2D). Alternatively, clines may be weaker or absent in the invasive range (Fig. 1E-G) due to weak selection across a narrow thermal gradient or evolutionary constraints imposed by bottlenecks.

## Materials and Methods

### Plant Propagation

In June 2019, we grew seeds from 18 North American perennial and 13 United Kingdom populations of *Mimulus guttatus* (Fig. 2A,B) in the North Carolina State University Phytotron. We used an average of 3 seed families from populations spanning a broad latitudinal and thermal gradient in the native and invasive ranges (see “Supplementary methods” for information regarding population selection), totaling 95 unique genotypes (Table S1). We sowed 3-5 seeds from each seed family into 72-cell plug trays filled with Fafard 4P soil (Fafard® 4P Mix, Sun Gro Horticulture) and topped with germination mix (Sunshine® Redi-Earth Plug & Seedling, Sun Gro Horticulture), placed them in a Percival LT-105X chamber, and subjected them to a 4°C cold, dark treatment with daily misting for one week. After one week, we transferred seedlings to a growth room, with a 16-hour day and 8-hour night photoperiod and a 20/15°C day/night temperature regime. Cold treatments were repeated for seed families with poor initial germination, which were primarily from the invasive range. After cold treatments, replanted families remained in the Percival chamber with 20°C days and colder (5°C) nights for two weeks before being moved to the growth room. Throughout seedling establishment and the remainder of the experiment, we watered plants in the growth room with a nutrient solution (NPK + micronutrients), dumped stagnant water, and randomized trays to reduce location effects twice per week. Three weeks after planting, seedlings were thinned to one, central-most individual per cell. Six weeks after planting, individuals were repotted into 8-inch pots with Fafard 4P. These large plants served as the source of replicate cuttings for thermal performance experiments and remained in the growth room for the duration of the experiment. We randomized the pots into new trays, each of which held 2 plants. Eight weeks after planting, we cut the primary stem of each plant to its base to encourage branching, which is ideal for taking replicate cuttings.

### Thermal Performance Experiment

Ten weeks after planting, plants possessed sufficient vegetative growth to support many cuttings. To generate thermal performance data for each genotype within each population, we took cuttings of similar sizes from these plants and grew them in 2.5-inch pots within 32-pot trays. As perennial *M. guttatus* often reproduces clonally in the native and invasive range (Truscott et al., 2006; van Kleunen, 2007; Pantoja et al., 2018), clones permit performance measurements of the same genotypes across a range of temperatures. Another advantage of using cuttings rather than seedlings is that they are likely less prone to maternal effects (reviewed in Roach & Wulff, 1987). Prior to temperature treatments, we allowed each set of clones 2 weeks to establish roots. During this period, cuttings were kept within chambers set to 20°C day/15°C night and bottom-watered daily with a nutrient solution. We randomized cuttings both within and among chambers twice per week to reduce location effects. We subsequently transferred clones into a growth chamber programmed to one of six temperature regimes. Over the course of the experiment, we randomly assigned six temperature treatments to one of three identical Percival LT-105X chambers (Percival Scientific, Inc.; Perry, Iowa, USA), resulting in two full rounds of chamber use. For each temperature regime, we took two replicates (clones) of each genotype and randomized them among six trays. We subjected clones to 16-hour days and 8-hour nights, according to procedures previously implemented by Paul et al. (2011), Sheth & Angert (2014), and Wooliver et al. (2020), with one of the following day/night temperature regimes (°C): 10/0, 20/10, 25/15, 30/20, 40/30, and 45/35. Temperature regimes were replicated once, along with an additional replicate of the 45°C/35°C regime. In total, we took 1,338 cuttings. However, 86 cuttings failed to establish, resulting in a final dataset of 1,252 individual clones from 18 native and 13 invasive populations, each comprising 54 and 41 genotypes, respectively. We exposed clones to a given temperature regime for one week, during which trays were sub-irrigated daily with water to prevent different rates of nutrient uptake at different temperatures.

We measured relative change in stem length over the 7-day temperature treatment. We chose this short treatment period because previous work with *M. guttatus* has documented significant growth responses to temperature in one week (Sheth & Angert, 2014). We quantified the length of the primary stem and the total length of branches (including stolons and side branches) before and after each temperature treatment (*stem_in_* and *stem_out_*, respectively). Total length of branches was estimated as the number of branches multiplied by the length of an average branch approximately halfway down the primary stem. We then calculated relative growth rate (RGR) as a proxy for fitness as the change in total stem length per initial stem length per day (Eq. 1). Negative RGR values arising when clones lost stem length or branches due to dieback at temperature extremes were set to zero, and we excluded negative RGR values resulting from accidental damage. Although RGR is not a measure of total reproductive output, we used this metric as a proxy for fitness because size has been related to reproductive output in other *Mimulus* species (Sheth & Angert, 2018).

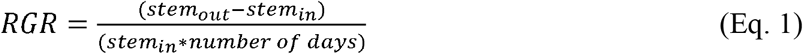

### Bayesian TPC Models

We performed all analyses in R v. 3.6.2 (R Core Team, 2019). To generate TPCs for native and invasive populations of *M. guttatus*, we used a hierarchical Bayesian model (R package performr v0.2; Tittes et al., 2019) to simultaneously fit curves predicting RGR across temperature for all populations while estimating uncertainty. As this model does not allow inclusion of random effects, we first averaged total RGR across clones of each genotype at each temperature. This model also required scaling RGR values by the mean RGR across all data and centering daytime temperature around zero. Thus, we rescaled model outputs to reflect actual RGR values and temperatures. To improve the effectiveness of posterior sampling, we altered the default settings of the model to include a total of 4,000 iterations per chain and a maximum tree depth of 15 (Gelman et al., 2014). This model utilizes a derivation of Kumaraswamy’s probability density function to fit performance curves across a continuous environmental gradient. We assessed the fit of Bayesian models using a posterior predictive *P*-value, which uses a test variable to give the probability that values drawn from the simulated posterior predictive distribution will exceed the observed values. *P*-values closest to 0.5 indicate adequate fit between the modeled and observed data (Gelman et al., 2014). The Bayesian *P*-value for the overall model was 0.53, and *P*-values for each population ranged from 0.2-0.82 (Table S2).

From these models, we obtained estimates of thermal optimum (*T_opt_*), thermal performance breadth (T_*breadth*_), maximum performance (*P_max_*), and area under the curve (*AUC*) for each population (Table S3). We selected critical values for *T_breadth_* from 100 equally spaced points along the temperature axis closest to the temperatures corresponding to 50% of *P_max_*. Although these models also generated estimates of critical upper and critical lower thermal limits (temperatures at which RGR decreases to zero), these estimates extended beyond our temperature treatments. We thus excluded these parameters from our analyses.

### Statistical Analyses for Hypothesis Testing

We tested for divergence in TPC parameters between native and invasive populations of *M. guttatus* independent of latitude by conducting post-hoc comparisons of TPC model iterations. For all iterations, we calculated the mean and 95% credible interval of the pairwise difference in each TPC parameter between the native and invasive range. We interpreted a statistically significant difference if the 95% credible interval did not include zero.

We examined the potential for specialist-generalist tradeoffs within either range using a general linear model with *T_breadth_*, range (native or invasive), and their interaction as predictors and *P_max_* as the response. We also ran a reduced linear model which removed the interaction term to account for the possibility of similar tradeoffs in both ranges. We used the Akaike information criterion (AIC) to assess model fit. For all linear models, we evaluated statistical significance using α=0.05. A negative relationship between *T_breadth_* and *P_max_* would indicate a specialist-generalist tradeoff, whereas a positive relationship would be consistent with a “master-of-all” scenario. Given a final model with a significant interaction term, we ran separate models for each range. To determine whether the assumption that the area under TPCs remained constant was supported, we conducted pairwise comparisons of population-level mean *AUC* based on model iterations. We used a 95% credible interval to assess significant differences between populations within each range and between populations of different ranges.

To determine whether TPCs of invasive populations have rapidly adapted to temperature and latitudinal gradients in the novel range, we compared latitudinal and thermal clines of TPC parameters in the native and invasive range using general linear models. To test whether *T_opt_* decreased with latitude or increased with mean annual temperature, we modeled *T_opt_* as a function of either latitude or mean annual temperature, range, and their interaction. Similarly, to evaluate whether *T_breadth_* increased with latitude in the native range, decreased with latitude in the invasive range, and increased with temperature seasonality in both ranges, we modeled *T_breadth_* as a function of latitude or temperature seasonality, range, and their interaction. We used AIC to compare models with and without the interaction term to account for the possibility of clines of similar magnitude and direction in both ranges. If the final model for each TPC parameter included a significant interaction term, we conducted range-specific models with only latitude or temperature as predictors. Shallower or nonsignificant slopes within the invasive range relative to the native range would suggest a lack of parallel evolution in response to thermal gradients. In contrast, a significant effect of latitude or temperature, along with a nonsignificant or absent interaction term would indicate parallel evolution of clines across ranges, though we note that a nonsignificant interaction term could also arise due to a lack of statistical power.

## Results

### Overall Shifts in TPC Parameters Between the Native and Invasive Ranges

We found no support for the hypothesis that *M. guttatus* has evolved in response to novel thermal regimes within its invasive range via TPC shifts. Pairwise comparisons revealed that *T_opt_* and *T_breadth_* exhibited 95% credible intervals that included zero (pairwise difference for *T_opt_*= −0.654°C, 95% CI= −1.405, 0.108; pairwise difference for *T_breadth_*=0.018°C, 95% CI= −1.191, 1.239), indicating that neither differed significantly between ranges (Fig. 3, Fig. S3). Additionally, pairwise comparisons showed that *AUC* and *P_max_* did not differ between ranges (pairwise difference for *AUC*= −3.043, 95% CI= −7.485, 1.344; pairwise difference for *P_max_* = −0.071 cm/cm/day, 95% CI= −0.194, 0.046; Fig. S3).

**Figure 3.**
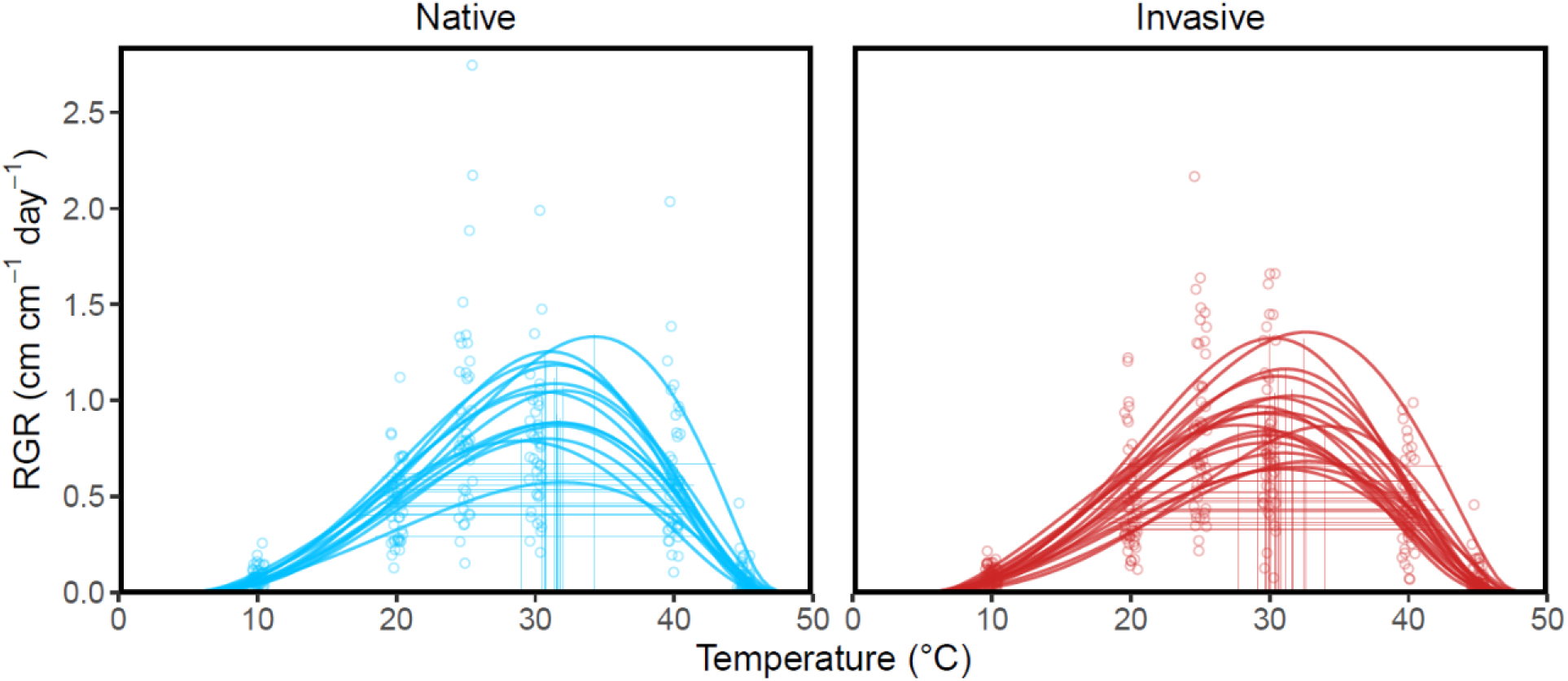
Thermal performance curves of 18 native and 13 invasive populations of *M. guttatus*. Vertical lines represent thermal optima (*T_opt_*) and horizontal lines denote thermal performance breadth (*T_breadth_*). Points represent genotype-level mean relative growth rate (RGR) at a given daytime temperature.

### Specialist-Generalist Tradeoffs

Consistent with specialist-generalist tradeoffs, *P_max_* decreased with *T_breadth_* (Table 1, Fig. 4). The final model included an interaction between *T_breadth_* and range (Table 1). However, this interaction was not statistically significant, suggesting that the direction and strength of the relationship between *T_breadth_* and *P_max_* did not differ between the native and invasive ranges (Table 1). The main effect of range was also non-significant (Table 1). Failing to meet the assumption that area is constant when there are specialist-generalist tradeoffs, pairwise comparisons indicate that *AUC* significantly differed among populations both within and between ranges (Palaima & Spitze, 2004; Fig. S2).

**Table 1.** Regression coefficients and *P*-values from full general linear models relating response variables to predictors. β1 indicates the main effect of either latitude, mean annual temperature (MAT), thermal performance breadth (*T_breadth_*), temperature seasonality (TS), or performance maximum (*P_max_).* β2 indicates the main effect of range (invasive vs. native). β3 indicates the interaction between range and the predictor corresponding to β1. Based on AIC, we chose bolded models as the final model for each respective cline.

| Response | Predictor ($\beta 1, \beta 2, \beta 3$) | Full Model | | | AIC | Adj. $R^2$ |
| --- | --- | --- | --- | --- | --- | --- |
| | | $\beta 1, P$ | $\beta 2, P$ | $\beta 3, P$ | | |
| $P_{max}$ | **$T_{breadth}, R, T_{breadth} * Range$** | **-0.140, $P=0.008^{**}$** | **-1.963, $P=0.146$** | **0.088, $P=0.159$** | **-10.448** | **0.219** |
| | $T_{breadth}, R$ | -0.083, $P=0.009^{**}$ | -0.070, $P=0.323$ | --- | -10.131 | 0.188 |
| $T_{opt}$ | Lat, R, Lat*R | 0.045, $P=0.712$ | 3.812, $P=0.590$ | -0.087, $P=0.504$ | 111.848 | -0.002 |
|  | **Lat, R** | **-0.033, $P=0.409$** | **-0.908, $P=0.123$** | --- | **110.372** | **0.018** |
| | MAT, R, MAT*R | 0.038, $P=0.899$ | -1.738, $P=0.521$ | 0.115, $P=0.710$ | 107.545 | 0.128 |
|  | **MAT, R** | **0.148, $P=0.028^*$** | **-0.748, $P=0.108$** | --- | **105.708** | **0.155** |
| $T_{breadth}$ | **Lat, R, Lat\*R** | **-0.201, $P=0.069$** | **-11.209, $P=0.077$** | **0.204, $P=0.079$** | **103.385** | **0.020** |
| | Lat, R | -0.018, $P=0.623$ | -0.120, $P=0.820$ | --- | 104.993 | -0.062 |
| | TS, R, TS*R | 0.058, $P=0.407$ | 1.265, $P=0.689$ | -0.034, $P=0.640$ | 104.236 | -0.008 |
|  | **TS, R** | **0.026, $P=0.117$** | **-0.197, $P=0.662$** | --- | **102.492** | **0.020** |

$^*=P<0.05$ , $^{**}=P<0.01$ , $^{***}=P<0.001$

**Figure 4.**
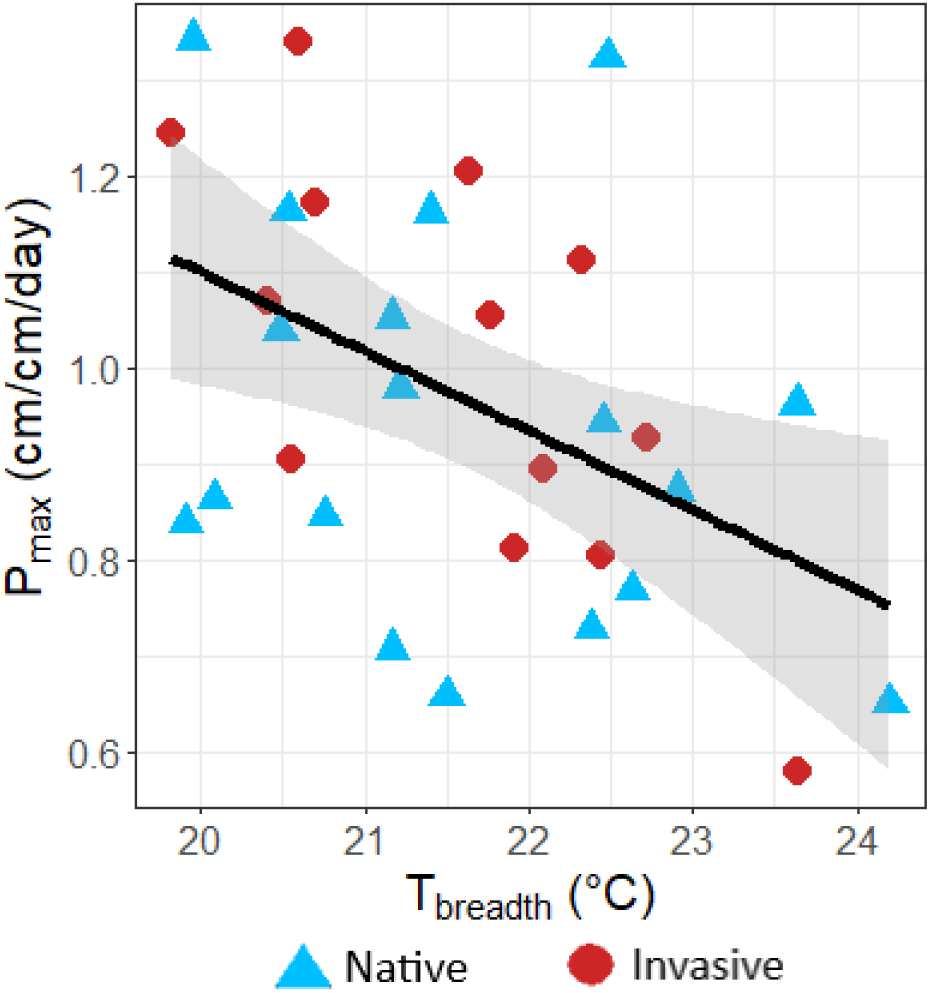
Specialist-generalist tradeoffs as shown by a relationship between maximum performance (*P_max_*) and thermal performance breadth (*T_breadth_*) across populations of *M. guttatus*. Black line indicates the main effect of *T_breadth_* (α=0.05; Table 1) from the full model with the interaction term removed, and grey shading shows 95% confidence interval.

### Clines of TPC Parameters

Overall, latitudinal and thermal clines varied in strength depending on the TPC parameter and/or the geographic range. The final models for *T_opt_* as a function of latitude or mean annual temperature (MAT) did not include an interaction between latitude or MAT and range (Table 1). Failing to support our hypothesis, *T_opt_* did not decrease with latitude (Table 1, Fig. 5A). Further, there was no main effect of range when assessing clines of *T_opt_* across latitude (Table 1). *T_opt_* increased with MAT, but there was also no main effect of range on *T_opt_* when assessing clines across MAT (Table 1, Fig. 5B). To further dissect the relatively low variance explained by this model (Adj. *R^2^*=0.155), we conducted range-level models. We found that the cline of *T_opt_* increasing with MAT in our full model was primarily driven by adaptive differentiation within the native range (β=0.153, *P=*0.036, Adj. *R^2^*=0.199), whereas there was no relationship between MAT and *T_opt_* in the invasive range (β=0.038, *P=*0.899, Adj. *R^2^*=−0.089).

**Figure 5.**
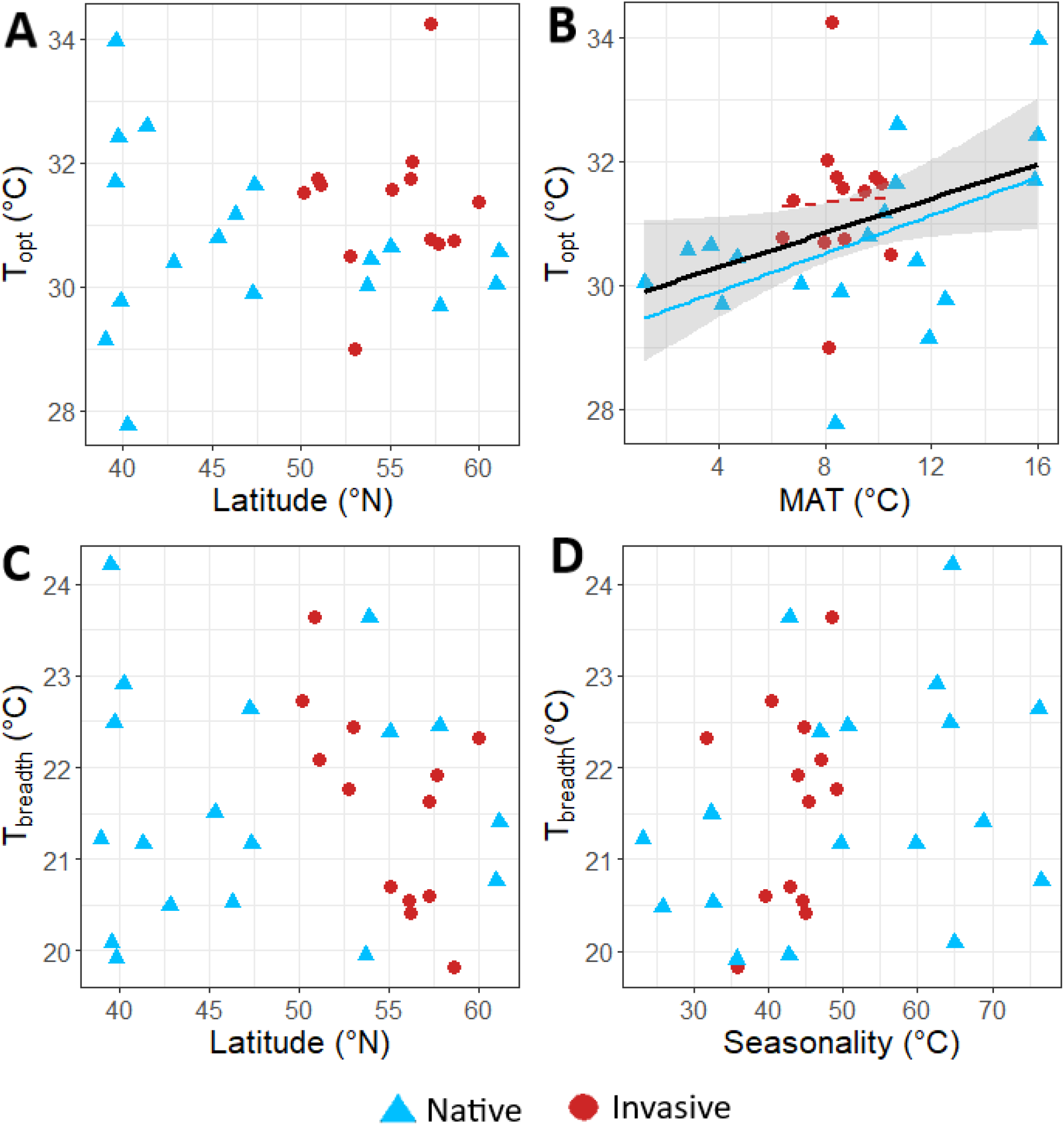
Relationships between population-level thermal optimum (*T_opt_*) or thermal performance breadth (*T_breadth_*) and latitude, mean annual temperature (MAT), or temperature seasonality. A main effect regression line (black) is shown to indicate an overall cline of MAT (α=0.05; Table 1) with *T_opt_*. However, this cline was driven by significant adaptive differentiation within the native range (solid blue line; *P*<0.05), rather than the invasive range (dashed red line; *P*>0.05).

The final model of *T_breadth_* as a function of latitude included an interaction between latitude and range (Table 1). Contrary to the prediction that *T_breadth_* would increase with latitude in the native range and decrease with latitude in the invasive range, *T_breadth_* showed no relationship with latitude (Table 1, Fig. 5C). There was not a significant main effect of range on *T_breadth_*, and the relationship between latitude and *T_breadth_* did not differ between ranges (Table 1). The final model of *T_breadth_* as a function of temperature seasonality did not include an interaction term (Table 1). We found no cline of *T_breadth_* with temperature seasonality (Table 1; Fig. 5D), failing to support the climate variability hypothesis. Further, there was no main effect of range on *T_breadth_* when assessing clines of *T_breadth_* across temperature seasonality (Table 1).

## Discussion

We compared thermal performance curves (TPCs) of 18 native and 13 invasive populations of *Mimulus guttatus* to test key hypotheses about the role of climatic niche evolution in facilitating biological invasions. First, our results provided no support for the hypothesis that thermal optimum and breadth vary between the native and invasive ranges (Fig. S3). Second, failing to support the hypothesis that *M. guttatus* is a “master-of-all,” there were specialist-generalist tradeoffs in both the native and invasive ranges (Table 1, Fig. 4). Finally, contrary to the hypothesis that native and invasive ranges show parallel phenotypic clines, there was limited evidence for parallel clines in TPC parameters between ranges (Table 1, Fig. 5). Rather than supporting the hypothesis that TPC evolution has played a role in facilitating *M. guttatus* invasion in the UK, these results provide physiological support for thermal niche conservatism in the invasive range. Below, we discuss the implications of these findings in light of the evolutionary processes that could contribute to biological invasion.

### Evolution of Thermal Performance Curves

We found no support for the hypothesis that invasive populations have undergone changes in mean TPC parameters relative to native populations (Fig. 3, Fig. S3). Instead, *T_opt_* and *T_breadth_* were similar between native and invasive ranges, consistent with the finding of climatic niche conservatism between the native North American and invasive European ranges of *M. guttatus* based on niche models (Da Re et al., 2020). However, like many other invasive species (Liu et al., 2020), the invasive range of *M. guttatus* has undergone substantial niche unfilling (Da Re et al., 2020), where invasive populations are found in a comparatively small subset of the climatic conditions (including temperature; Fig. S1) occupied in the native range. Although the range of temperatures is narrower in the introduced range (Fig. 2C), *T_breadth_* did not differ between ranges (Fig. S3), suggesting that invasive *M. guttatus* populations are poised to occupy greater climatic niche space should it become available. Maintenance of broad thermal tolerance in the invasive range may allow *M. guttatus* populations to withstand rapid changes in temperatures, increasing risks that invasive populations will proliferate under climate change. Future TPC comparisons that include introduced populations from additional regions would further contribute to our understanding of the role of thermal niche evolution in the global range expansion of *M. guttatus*.

One factor that could contribute to the evolutionary stasis of TPC parameters in the invasive range is insufficient standing genetic variation (Prentis et al., 2008; Dlugosch et al., 2015; Colautti et al., 2017). Although lower neutral genetic variation in the introduced range (Puzey & Vallejo-Marín, 2014; Pantoja et al., 2017) could have constrained the rapid evolution of *T_opt_* and *T_breadth_* in *M. guttatus*, the highly admixed nature of invasive populations (Vallejo-Marín et al., 2020) make this possibility unlikely. Instead, genetic variation resulting from admixed invasive populations likely contributed to the maintenance of broad TPCs and allowed these populations to easily tolerate the narrower range of temperatures in the UK. In sum, the invasive populations have yet to undergo significant divergence in TPCs from native populations due to overlap in thermal niche space, rather than lacking the genetic variation necessary for evolution to occur. If invasive populations exhibit high admixture, they may spread further via thermal niche evolution in the face of climate change.

### Specialist-Generalist Tradeoffs

We found that TPCs in both ranges were constrained by specialist-generalist tradeoffs, such that broad TPCs came at the cost of lower maximum performance (Table 1, Fig. 4). Considering the similar *T_breadth_* across ranges, maintenance of broad thermal tolerance in the narrow thermal conditions of the invasive range may incur significant performance costs via specialist-generalist tradeoffs. However, variation in *AUC* among populations within each range (Fig. S2) suggests that the assumption that area remains constant does not hold for *M. guttatus* (Palaima & Spitze, 2004). As such, the tradeoff between *T_breadth_* and *P_max_* is only partial. Although *AUC* and *T_breadth_* were not correlated (ρ=−0.2692, *P*= 0.1431), suggesting that thermal niche space is not affected by variance in *T_breadth_*, *AUC* was positively correlated with *P_max_*, (ρ=0.9767, *P*<0.001). These results imply that variation in TPCs among populations is partly explained by vertical shifts across a fast-slow spectrum of performance (Izem & Kingsolver, 2005), where some populations achieve a greater *AUC* by having both a high maximum performance and a broad TPC (Palaima & Spitze, 2004). Thus, variation in TPCs in native and invasive populations of *M. guttatus* seems to be controlled by both vertical shifts where high-performing populations are ‘masters-of-all’ and specialist-generalist tradeoffs where generalist populations are ‘masters-of-none’.

### Evolution of Clines

We hypothesized that phenotypic clines exhibited in the native range would be repeated in the invasive range. Although temperature varies predictably with latitude in the invasive range (Fig. 2C,D), and there is ample evidence that rapid adaptation can facilitate invasion success (Oduor et al., 2016), our results did not support the hypothesis of parallel clines. Rather, *T_opt_* increased with mean annual temperature across both ranges, though the amount of variance explained was modest (*Adj. R^2^*=0.155). As native populations were the primary driver of the cline detected in the full model (Fig. 5B), adaptive differentiation of *T_opt_* to mean annual temperature may contribute to the success of this species in its native range. The absence of this cline in the invasive range implies weaker selection on *T_opt_* or insufficient time for adaptation in the invasive range. Given that invasive populations of *M. guttatus* have maintained similar *T_breadth_* as native populations, selection in the invasive range with a relatively narrow range of mean annual temperatures would not likely favor adaptive differentiation in *T_opt_* (Fig. 2C).

Failing to support the climate variability hypothesis (Dobzhansky, 1950; Janzen, 1967; Stevens, 1989; Gutiérrez-Pesquera et al., 2016), *T_breadth_* was not related to latitude or temperature seasonality in either range. There are many possible explanations for a lack of clines in *T_breadth_* in native populations. First, since six native populations were inland perennials, differing regimes of temperature seasonality in coastal and inland environments of perennial *M. guttatus* could mask latitudinal and macroclimatic trends in *T_breadth_* (Hall & Willis, 2006; Lowry et al., 2009). Second, microclimatic variation could cause populations of *M. guttatus* to occupy areas with either highly similar or drastically different regimes of temperature seasonality, which could hinder the detection of clines at broader macroclimatic scales (Franco & Nobel, 2003; Rashkovetsky et al., 2006; De Frenne et al., 2013). While additional analyses of variation in TPC parameters within populations would be helpful for dissecting this hypothesis, our model was unable to fit genotype-level curves due to insufficient replication. Finally, high gene flow among native populations could swamp adaptation of TPCs to local thermal regimes (Paul et al., 2011). Rather than evolving TPCs in response to thermal variation, native populations may consist of general-purpose genotypes with wide TPCs which contributed to their successful establishment as an invasive species (van Kleunen et al., 2011). This conclusion is consistent with the finding of high rates of gene flow among coastal populations occupying similar latitudes in the native range relative to our study populations (Twyford & Friedman, 2015). Nonetheless, our findings are particularly surprising given the evolution of parallel phenotypic clines in the introduced ranges of several species (van Boheemen et al., 2019; Hernández et al., 2019; McGoey et al., 2020). Overall, our results suggest that though native *M. guttatus* populations are adaptively differentiated by thermal optima, populations are equally tolerant across a broad temperature gradient in western North America. Such homogenization of *T_breadth_* among native populations may have allowed *M. guttatus* to maintain a broad thermal niche space upon introduction.

### Caveats

There are many caveats which may have impacted the results of our study. First, our temperature regimes maintained constant respective day and night temperatures, but in nature plants experience temperature fluctuations. Second, although we incorporated multiple measurements of stem and branch growth, populations may vary in their investment in stem and branch growth versus other growth parameters (such as leaf number or belowground growth) at different temperatures. Third, we focused on one performance metric, RGR over a week-long period, but to fully understand performance tradeoffs and variation in performance across temperatures, multiple performance metrics (e.g., survival and fecundity) should be measured over longer experimental periods and across multiple life stages. Ultimately, performance metrics that better approximate lifetime fitness could yield stronger phenotypic clines. Nevertheless, short-term RGR of cuttings captures performance of clonally reproducing plants characteristic of perennial *M. guttatus*. Finally, we focused on temperature, one axis of the climatic niche, but evolution along other niche axes such as precipitation and edaphic properties could also contribute to invasion success (Hall & Willis, 2006; van Kleunen & Fischer, 2008). Further, the biotic environment may also differ in the introduced range (Holeski et al., 2013; Thawley et al., 2018), implying that many factors beyond abiotic conditions can play a part in adaptation to novel conditions during invasions.

## Conclusions

Plant invasions have been widely studied for land management purposes, but little consensus exists on the ecological and evolutionary processes that facilitate invasion success. Our findings that thermal optima and breadth are similar in the invasive and native ranges, and that populations did not exhibit clines in thermal breadth even in the native range, suggest that general-purpose genotypes, rather than rapid TPC evolution, may have contributed toward successful establishment in the invasive range. Our results have important implications toward discovering what traits are favored for invasive organisms. For example, although having a broad thermal performance curve may aid in initial establishment of populations in a new environment, specialist-generalist tradeoffs may constrain the further evolution of broader TPCs. By comparing the evolution of thermal performance across broad environmental gradients in a species native and invasive range, we can predict which species hold the potential to become aggressive invaders and forecast how they may fare under rapid climate change. Species that are genetically predisposed towards a broad thermal tolerance may rapidly expand in their invasive range as climatic conditions shift. These predictions will be invaluable in preparing for future species invasions, strengthening our efforts to manage invasive species in the face of rapid climate change and globalization.

## Acknowledgements

We thank B. Caldwell, E. Coughlin, N. Gold, M. Kidd, D. Ryan, J. Torres, E. Vtipil, and M. Wiegmann for assistance with data collection and plant care. We also thank NCSU Phytotron staff members for assistance with plant care and chamber maintenance. We are grateful to B. Blackman, J. Colicchio, J. Coughlan, J. Friedman, L. Holeski, D. Lowry, and M. Rotter for generously providing seeds for this work. W. Hoffmann, M. Burford-Reiskind, Associate Editor Peter Alpert, and two anonymous reviewers provided valuable feedback on earlier drafts, and M. Flores-Vergara provided guidance about plant propagation. This project was funded by USDA National Institute of Food and Agriculture Hatch 1016272.

## Author Contributions

A.Q. and S.N.S. conceived of and designed the study with feedback from M.V.M. A.Q. collected the data. A.Q. and R.W. analyzed the data. A.Q. and S.N.S. led the writing, with substantial contributions from R.W. and M.V.M. All authors contributed critically to the drafts and gave final approval for publication.

## Data Availability

All data and scripts associated with this study are available at https://github.com/akquerns/Guttatus_EvolTPCClines.

