## Supporting Information for "The evolution of thermal performance in native and invasive populations of *Mimulus guttatus*"

**Supplementary Methods, Tables and Figures**

**Supplementary methods:** Supplementary methods describing selection criteria for populations used in thermal performance experiments

To compare thermal performance curves of native and invasive populations of *M. guttatus*, we propagated field-collected seeds from 1-15 families in each of 32 perennial populations from the native North American range and 20 perennial populations from the invasive United Kingdom range (Table S1), totaling 346 seed families (see “Propagation of Mother Plants” below). To select focal populations among the populations that successfully germinated (26 native and 13 invasive populations), we conducted a principal component analysis (PCA). We used the geospatial coordinates of these populations to extract data on 11 temperature variables (1970-2000, ~1km resolution) from WorldClim v. 2 (Fick & Hijmans, 2017) for all populations in which at least one seed family germinated (Table S1). We used these variables to conduct a PCA to understand how climate varies within and between the native and invasive ranges (Fig. S1).

The PCA indicated two principal components explaining the majority of variance across populations (Fig. S1). The first principal component, which explained 53.94% of variance, was characterized primarily by differences in mean temperature of the warmest quarter, mean temperature of the driest quarter, and mean annual temperature. The second principal component, which explained 32.74% of variance, was characterized by differences in maximum temperature of the warmest month, minimum temperature of the coldest month, and annual temperature range. The first two principal components showed that invasive populations encompassed a small subset of temperature conditions occupied by native populations (Fig. S1).

Using PCA coordinates generated for each population, we selected native populations for our experiment which were distributed throughout the full range of temperatures in the native range without over-representation of specific climates. Specifically, over-representation was indicated by a population being within 0.2 units of another population on either PC axis. Some exceptions were made to include a higher number of Alaskan populations, which may be the source of UK invasive populations (Pantoja et al., 2017). From this analysis, we identified 18 populations that represented a wide breadth of temperatures within the native range (Fig. 2A). Due to lower germination success of genotypes from the invasive range in the UK, we used all 13 available populations (Fig. 2B). Within each population, we randomly selected an average of 3 seed families to represent in our experiment, representing a total of 95 unique genotypes from 31 populations (Table S1). During this process, plants exhibiting annual characteristics (diminutive size, lack of side branches and stolons, small flowers; Hall & Willis, 2006) were excluded in favor of perennial plants.

**Table S1.** Seed collection sites for potential *M. guttatus* populations to be included in thermal performance experiments. “Donor” describes the labs or individual collectors who donated seeds from each population. “Reference Population” describes the original population code or name provided by donors for each geographical occurrence. “New Population Code” describes the codes assigned to populations used in the full experiment, where N1-18 are native populations, I1-13 are invasive populations, and dashes indicate populations which were not used. “Number of Genotypes” describes the total number of seed families within each population which were donated for our experiment. “Genotypes Selected” describes the number of seed families within each population used in thermal performance experiments, given germination success and population selection criteria. “Range” describes whether the population exists within the North American (native) range or United Kingdom (invasive) range of *M. guttatus*.

| ***Donor*** | ***Reference Population*** | ***New Population Code*** | ***Latitude*** | ***Longitude*** | ***Number of Genotypes*** | ***Genotypes Selected*** | ***Range*** |
| --- | --- | --- | --- | --- | --- | --- | --- |
| Lowry | FRA | N1 | 39.0083 | -123.69395 | 7 | 3 | North America |
| Coughlan | TFP | --- | 39.53903 | -121.58286 | 15 | 0 | North America |
| Coughlan | TBM1 | N2 | 39.55229 | -121.567 | 15 | 3 | North America |
| Coughlan | CCR | N3 | 39.60485 | -121.598 | 15 | 3 | North America |
| Coughlan | HWY 70 | --- | 39.6153 | -121.6 | 15 | 0 | North America |
| Lowry | ANR | --- | 39.73687 | -123.631 | 11 | 0 | North America |
| Coughlan | BLU | N4 | 39.7462 | -121.6728833 | 14 | 3 | North America |
| Coughlan | CNV | --- | 39.813717 | -121.57175 | 2 | 0 | North America |
| Lowry | LJA | N5 | 39.860529 | -123.902367 | 11 | 3 | North America |
| Coughlan | CH | --- | 40.1028 | -121.4993 | 15 | 0 | North America |
| Coughlan | CHG | N6 | 40.2476 | -121.447433 | 12 | 3 | North America |
| Lowry | SAL | N7 | 41.339663 | -123.388162 | 11 | 3 | North America |
| Blackman | HMP | --- | 42.690011 | -124.447772 | 5 | 0 | North America |
| Lowry | BLN | N8 | 42.83772 | -124.56 | 7 | 3 | North America |
| Colicchio | 7DEV | --- | 43.235261 | -124.392128 | 7 | 0 | North America |
| Lowry | BLY | --- | 43.989674 | -123.177453 | 11 | 0 | North America |
| Coliccho | HOB | --- | 44.143124 | -124.122314 | 2 | 0 | North America |
| Coliccho | NEP | --- | 44.26739 | -124.108563 | 2 | 0 | North America |
| Colicchio | YRR | --- | 44.309916 | -124.101597 | 3 | 0 | North America |
| Coliccho | NPB | --- | 44.623434 | -124.067267 | 10 | 0 | North America |
| Coliccho | QRV | --- | 44.677008 | -124.07399 | 4 | 0 | North America |
| Lowry | LOK | N9 | 45.3382533 | -123.9771333 | 12 | 4 | North America |
| Lowry | DIS | N10 | 46.3053 | -124.072 | 10 | 3 | North America |
| Holeski | Nowhere Ditch | N11 | 47.287683 | -117.962933 | 5 | 3 | North America |
| Lowry | HOC | N12 | 47.3854 | -123.147 | 11 | 3 | North America |
| Holeski/Rotter | TSG | N13 | 53.698133 | -132.526217 | 3 | 3 | North America |
| Friedman | 16-DHS | N14 | 53.90448 | -166.51068 | 5 | 3 | North America |
| Friedman | 16-AKC | N15 | 55.05603 | -162.32816 | 5 | 3 | North America |
| Friedman | 16-MFM | N16 | 57.79456 | -152.59483 | 5 | 3 | North America |
| Holeski/Rotter | Anchor River | --- | 59.741133 | -151.7475 | 5 | 0 | North America |
| Holeski/Rotter | Bird Point Creek | N17 | 60.95245 | -149.4112167 | 5 | 3 | North America |
| Holeski/Rotter | Crooked Creek | N18 | 61.13825 | -146.32465 | 5 | 2 | North America |
| Vallejo-Marín | CRO | I1 | 50.162 | -5.293 | 5 | 3 | United Kingdom |
| Vallejo-Marín | DEA | I2 | 50.904513 | -0.77971674 | 3 | 2 | United Kingdom |
| Vallejo-Marín | HOU | I3 | 51.09699 | -1.5084 | 5 | 3 | United Kingdom |
| Holeski/Rotter | Exford Bridge | --- | 51.133 | -3.64177 | 3 | 0 | United Kingdom |
| Vallejo-Marín | BRA | I4 | 52.768 | 1.297 | 5 | 3 | United Kingdom |
| Holeski/Rotter | St. Catherine Pasture | --- | 52.98935 | -3.466 | 3 | 0 | United Kingdom |
| Vallejo-Marín | CER | I5 | 53.00598 | -3.54927 | 4 | 3 | United Kingdom |
| Holeski/Rotter | River Nith Bridge | I6 | 55.0635 | -3.60888 | 3 | 1 | United Kingdom |
| Holeski/Rotter | River Ayre | --- | 55.4615 | -4.6257 | 3 | 0 | United Kingdom |
| Holeski/Rotter | Coldstream Bridge | --- | 55.6548 | -2.23938 | 3 | 0 | United Kingdom |
| Holeski/Rotter | John Muir Footpath | --- | 55.99497 | -2.55667 | 3 | 0 | United Kingdom |
| Holeski/Rotter | Balfron Mud Field | --- | 56.0653 | -4.39088 | 3 | 0 | United Kingdom |
| Vallejo-Marín | TIL | I7 | 56.1473 | -3.74477 | 5 | 4 | United Kingdom |
| Vallejo-Marín | DBL | I8 | 56.19654 | -3.96856 | 5 | 4 | United Kingdom |
| Vallejo-Marín | BAL | I9 | 57.23748 | -2.06385 | 5 | 4 | United Kingdom |
| Vallejo-Marín | TOM | I10 | 57.25499 | -3.36777 | 5 | 3 | United Kingdom |
| Vallejo-Marín | DAL | I11 | 57.682614 | -4.2652576 | 5 | 3 | United Kingdom |
| Holeski/Rotter | Loch Broom Hill | --- | 57.82897 | -5.06625 | 3 | 0 | United Kingdom |
| Vallejo-Marín | BKN | I12 | 58.575 | -4.767 | 5 | 4 | United Kingdom |
| Vallejo-Marín | NIN | I13 | 59.97777 | -1.30036 | 5 | 4 | United Kingdom |

**Table S2.** Posterior predictive *P*-values generated for each *M. guttatus* population’s thermal performance curve. *P*-values close to 0.5 indicate adequate fit between the modeled and observed data (Gelman et al. 2014). Invasive populations are indicated by I1-I13, and native populations are indicated by N1-N18 (Table S1).

| Population | Bayesian p-value |
| --- | --- |
| I1 | 0.30525 |
| I2 | 0.56975 |
| I3 | 0.198375 |
| I4 | 0.482125 |
| I5 | 0.565375 |
| I6 | 0.54275 |
| I7 | 0.638 |
| I8 | 0.5215 |
| I9 | 0.532125 |
| I10 | 0.711625 |
| I11 | 0.698 |
| I12 | 0.821375 |
| I13 | 0.457125 |
| N1 | 0.542375 |
| N2 | 0.426 |
| N3 | 0.671375 |
| N4 | 0.33275 |
| N5 | 0.694625 |
| N6 | 0.64025 |
| N7 | 0.2865 |
| N8 | 0.668625 |
| N9 | 0.3945 |
| N10 | 0.3935 |
| N11 | 0.644875 |
| N12 | 0.473875 |
| N13 | 0.565625 |
| N14 | 0.741375 |
| N15 | 0.493375 |
| N16 | 0.6755 |
| N17 | 0.50725 |
| N18 | 0.401875 |

**Table S3.** Mean thermal performance curve parameter values for each population, and 95% credible intervals for these parameter values, generated using a Bayesian model. Parameters are either in units of °C (*T_opt_* and *T_breadth_*), cm/cm/day (*P_max_*), or arbitrary units (*AUC*).

| Population | *T_opt_* | *T_breadth_* | *P_max_* | *AUC* |
| --- | --- | --- | --- | --- |
| I1 | 31.52 [29.21, 33.82] | 22.72 [18.49, 26.23] | 0.93 [0.67, 1.26] | 43.65 [31.88, 57.35] |
| I2 | 31.75 [29.06, 34.42] | 23.64 [19.09, 27.29] | 0.58 [0.44, 0.78] | 28.31 [21.66, 35.71] |
| I3 | 31.65 [29.72, 33.5] | 22.08 [19.1, 25] | 0.89 [0.7, 1.14] | 41.15 [33.39, 49.8] |
| I4 | 30.5 [28.93, 32.04] | 21.77 [19.28, 24.48] | 1.05 [0.84, 1.33] | 47.86 [39.89, 57.26] |
| I5 | 29.01 [26.43, 31.76] | 22.44 [18.71, 25.88] | 0.81 [0.59, 1.09] | 37.48 [27.85, 48.64] |
| I6 | 31.58 [28.98, 34.09] | 20.7 [16.94, 24.89] | 1.17 [0.69, 1.94] | 50.31 [32.05, 75.63] |
| I7 | 31.75 [29.96, 33.54] | 20.55 [17.82, 23.5] | 0.9 [0.68, 1.21] | 38.99 [30.4, 49.08] |
| I8 | 32.03 [30, 33.92] | 20.41 [16.98, 23.6] | 1.07 [0.83, 1.37] | 45.85 [36.12, 56.65] |
| I9 | 34.25 [32.71, 35.79] | 20.6 [17.91, 23.57] | 1.34 [0.98, 1.82] | 57.91 [44.47, 73.49] |
| I10 | 30.78 [28.62, 32.93] | 21.63 [18.68, 24.84] | 1.21 [0.88, 1.61] | 54.44 [41.5, 69.29] |
| I11 | 30.7 [28.5, 32.83] | 21.92 [18.56, 25.44] | 0.81 [0.59, 1.11] | 37.01 [28.47, 47.08] |
| I12 | 30.74 [28.51, 32.86] | 19.82 [16.75, 23.14] | 1.24 [0.87, 1.73] | 51.82 [38.57, 68.01] |
| I13 | 31.36 [29.32, 33.34] | 22.32 [19.54, 25.33] | 1.11 [0.83, 1.47] | 51.67 [40.3, 64.61] |
| N1 | 29.14 [26.94, 31.15] | 21.22 [18.16, 24.63] | 0.98 [0.68, 1.41] | 43.18 [32.11, 57.18] |
| N2 | 31.69 [29.39, 33.94] | 24.2 [20.77, 27.37] | 0.65 [0.51, 0.83] | 32.44 [26.22, 39.09] |
| N3 | 33.96 [31.47, 36.19] | 20.08 [16.15, 24.54] | 0.86 [0.55, 1.34] | 36.19 [24.8, 51.25] |
| N4 | 32.42 [30.39, 34.36] | 22.48 [19.61, 25.51] | 1.32 [0.99, 1.78] | 61.76 [48.7, 77.57] |
| N5 | 29.77 [27.58, 31.83] | 19.91 [16.79, 23.12] | 0.84 [0.64, 1.1] | 35.09 [27.86, 43.55] |
| N6 | 27.77 [25.39, 30.07] | 22.91 [19.84, 26.18] | 0.87 [0.61, 1.24] | 41.2 [30.34, 54.83] |
| N7 | 32.59 [30.31, 34.7] | 21.17 [17.17, 25.15] | 0.71 [0.5, 0.98] | 31.15 [23.02, 40.49] |
| N8 | 30.39 [28.56, 32.12] | 20.48 [17.76, 23.28] | 1.04 [0.8, 1.34] | 44.63 [35.83, 54.83] |
| N9 | 30.8 [28.54, 32.93] | 21.5 [17.77, 24.96] | 0.66 [0.51, 0.85] | 29.49 [23.66, 35.9] |
| N10 | 31.17 [29.09, 33.25] | 20.53 [17.46, 23.66] | 1.16 [0.85, 1.56] | 50.08 [37.76, 64.04] |
| N11 | 29.89 [28.04, 31.74] | 22.63 [19.81, 25.46] | 0.77 [0.62, 0.96] | 36.11 [30.13, 42.98] |
| N12 | 31.64 [29.9, 33.34] | 21.17 [18.09, 24.32] | 1.05 [0.76, 1.46] | 46.33 [35.52, 59.9] |
| N13 | 30.02 [27.45, 32.72] | 19.95 [16.64, 23.34] | 1.34 [0.94, 1.89] | 56.24 [41.07, 75.04] |
| N14 | 30.43 [28.08, 32.76] | 23.64 [20.36, 26.83] | 0.96 [0.69, 1.32] | 46.86 [34.46, 61.44] |
| N15 | 30.65 [27.74, 33.64] | 22.38 [17.64, 26.28] | 0.73 [0.54, 0.97] | 33.87 [24.91, 43.6] |
| N16 | 29.69 [27.21, 32.22] | 22.45 [18.96, 25.51] | 0.94 [0.72, 1.22] | 44.09 [33.42, 55.39] |
| N17 | 30.04 [27.28, 32.8] | 20.76 [16.83, 24.46] | 0.85 [0.62, 1.15] | 36.72 [27.09, 48.09] |
| N18 | 30.57 [28.34, 32.72] | 21.4 [18.2, 24.87] | 1.16 [0.81, 1.64] | 51.79 [37.61, 68.94] |

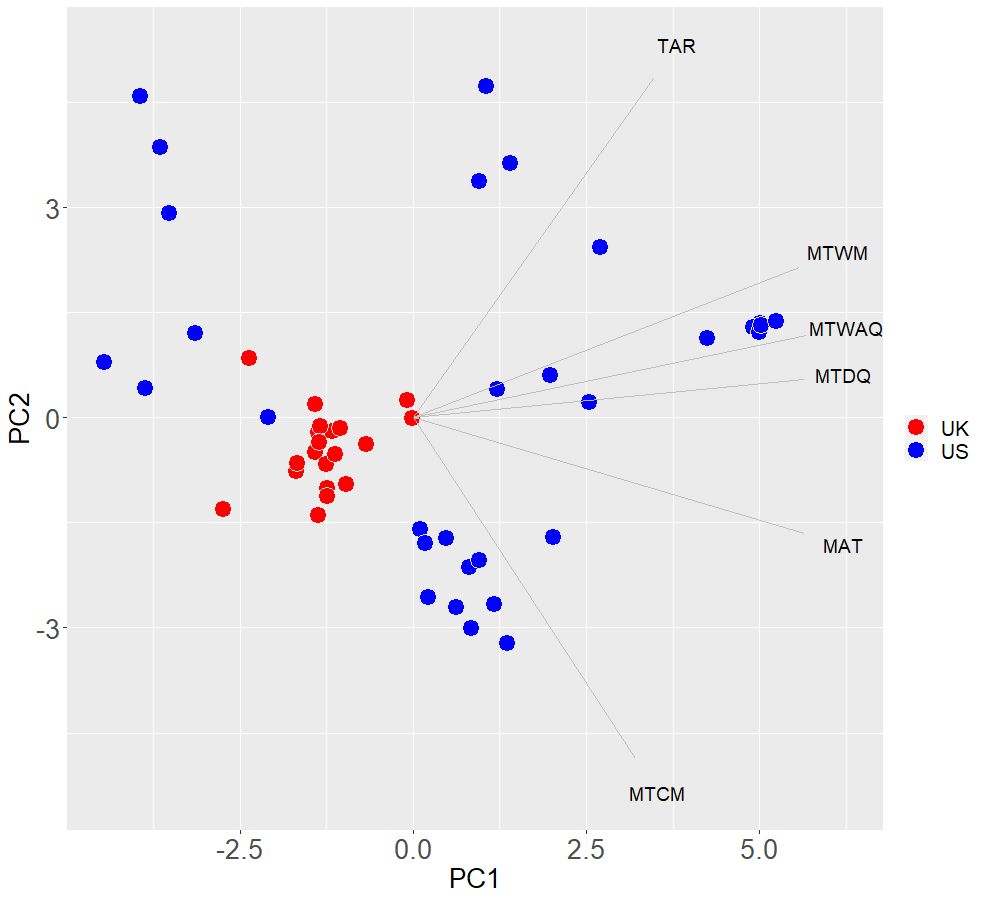

**Figure S1.** Results of PCA exploring variation in 11 temperature variables from WorldClim v. 2.0 across all populations considered for use in our study (Supplementary methods, Table S1; 1970-2000, Fick & Hijmans 2017). These variables include mean annual temperature (MAT), mean diurnal range (MDR), isothermality (ISO), temperature seasonality (TS), maximum temperature of the warmest month (MTWM), minimum temperature of the coldest month (MTCM), temperature annual range (TAR), mean temperature of the wettest quarter (MTWQ), mean temperature of the driest quarter (MTDQ), mean temperature of the warmest quarter (MTWAQ), and mean temperature of the coldest quarter (MTCQ).

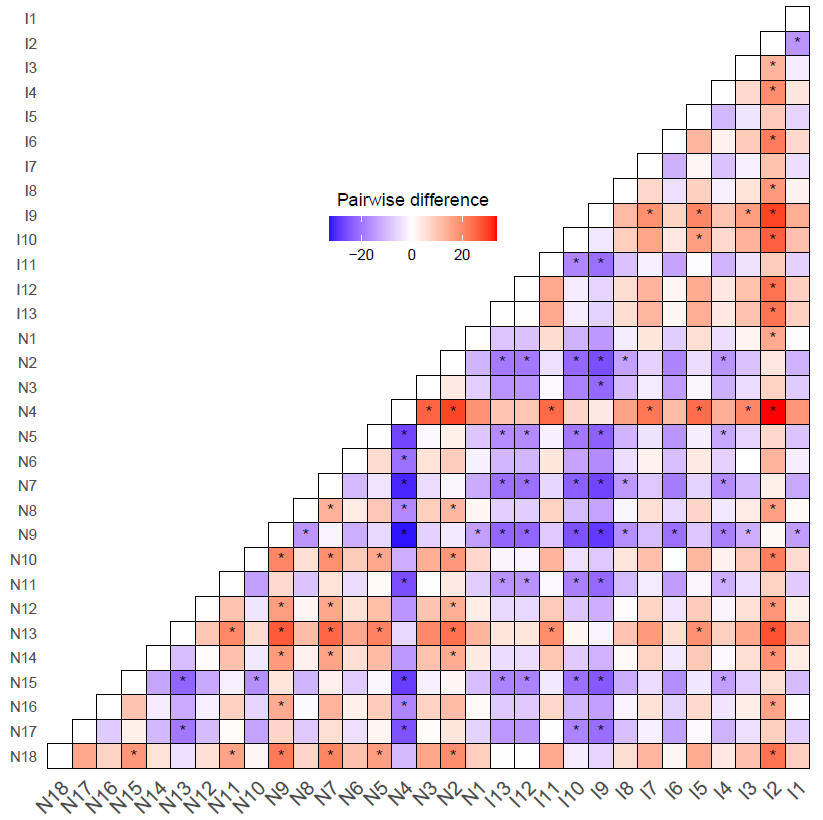

**Figure S2.** Pairwise comparisons of area under the thermal performance curve (AUC) across 31 invasive and native populations of *M. guttatus*. Population codes are in Table S1. The color of each box represents the mean pairwise difference between the x group (columns) and the y group (rows) for 8,000 iterations of a Bayesian model. Red indicates cases where the y group has a parameter value that is greater than the x group, and blue indicates cases where the y group has a parameter value that is less than the x group. Asterisks represent pairwise comparisons where the 95% credible interval for differences across all iterations of the model does not overlap zero.

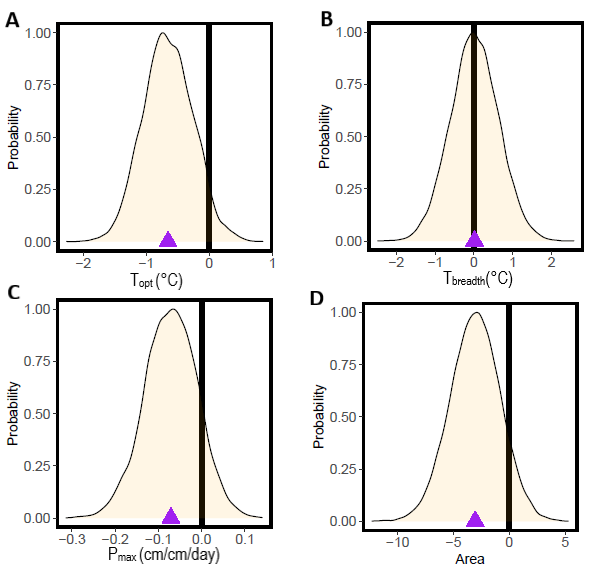

**Figure S3.** Pairwise comparisons of range-level mean thermal performance curve parameters. Pairwise differences are calculated by subtracting invasive range parameter estimates from native range parameter estimates. A positive difference indicates a higher parameter estimate in the native range, and a negative difference indicates a higher parameter estimate in the invasive range. Each bell curve shows a probability curve for pairwise differences based on 95% credibility. The mean pairwise difference is indicated by a purple triangle, and a vertical black line is shown to indicate a lack of difference in parameter estimates between the native and invasive range.
